# Activation of Histone 3 Lysine 9 methyl writing and reading capabilities within the G9a-GLP heterodimer

**DOI:** 10.1101/2021.05.05.442822

**Authors:** Nicholas A Sanchez, Lena M Kallweit, Michael J Trnka, Charles L Clemmer, Bassem Al-Sady

## Abstract

Unique among metazoan repressive histone methyltransferases, G9a and GLP, which chiefly target histone 3 lysine 9 (H3K9), require dimerization for productive H3K9 mono (me1)- and dimethylation (me2) *in vivo*. Intriguingly, even though each enzyme can independently methylate H3K9, the predominant active form *in vivo* is a heterodimer of G9a and GLP. How dimerization influences the central H3K9 methyl binding (“reading”) and deposition (“writing”) activity of G9a and GLP, and why heterodimerization is essential *in vivo* remains opaque. Here, we examine the H3K9me “reading” and “writing” activities of defined, recombinantly produced homo- and heterodimers of G9a and GLP. We find that both reading and writing are significantly enhanced in the heterodimer. Compared to the homodimers, the heterodimer has higher recognition of H3K9me2, and a striking ∼ 10-fold increased turnover rate for nucleosomal substrates under multiple turnover conditions, which is not evident on histone tail peptide substrates. Crosslinking Mass Spectrometry suggests that differences between the homodimers and the unique activity of the heterodimer may be encoded in altered ground state conformations, as each dimer displays different domain contacts. Our results indicate that heterodimerization may be required to relieve autoinhibition of H3K9me reading and chromatin methylation evident in G9a and GLP homodimers. Relieving this inhibition may be particularly important in early differentiation when large tracts of H3K9me2 are deposited by G9a-GLP, which may require a more active form of the enzyme.

## Introduction

The animal genome is partitioned into active and inactive regions by gene-repressive structures such as heterochromatin that restrict access to gene-activating factors (1). In early mammalian development, the genome of embryonic stem cells is highly transcriptionally active and features little heterochromatin marked by histone 3 lysine 9 methylation (H3K9me) (2). As differentiation begins, H3K9me heterochromatin expands, remodeling the genome and restricting fate (3). These expansions are carried out by the SETDB1 H3K9 trimethylase (me3) (3, 4) and the G9a and GLP H3K9 mono-and dimethylases (me1/me2). G9a and GLP direct the expansion of large tracks of H3K9me2, which can adopt a lineage-specific pattern (5). This heterochromatin expansion represses lineage inappropriate genes directly or silences enhancers that in turn direct several genes (6).

Like several heterochromatic methyltransferases, G9a and GLP have the capacity to both read and write H3K9me (7–10). Both enzymes contain an Ankyrin (ANK) repeat domain, which confers methyl-histone binding (reading) activity, and a SET domain, which confers methyltransferase (writer) activity. The presence of methylated substrates appears to stimulate G9a or GLP catalytic activity in an ANK domain-dependent manner (10). Other H3K9 methylases require such positive feedback for either lateral spreading (11, 12) or epigenetic maintenance (13, 14). How the ANK and SET domains of G9a and GLP regulate one another is opaque but may be central to understanding their function in cell fate control.

Unlike other metazoan H3K9 methyltransferases, G9a and GLP must associate with each other to carry out H3K9 methylation (9): When the interaction between the two enzymes is broken in vivo, the bulk of H3K9me1 and me2 is lost (15). This is despite the observation that each enzyme is individually capable of methylating histones in vitro (8–10, 16, 17). We hypothesized that forming the G9a-GLP complex (G9a-GLP) has a direct regulatory effect on the H3K9 methylation reaction. However, the biochemical and biophysical nature of homo- and heterotypic associations between G9a and GLP are poorly understood, as are any effects on H3K9me writing and reading that these associations may have.

In this study, we investigated the regulation heterodimerization imposes on G9a and GLP’s ability to read and write H3K9me.

## Results

### G9a and GLP form stable dimers at 1:1 stoichiometry

To directly assay the interplay between reading, writing, and dimerization, we expressed and purified G9a and GLP truncated to the C-terminal ANK and SET domains (ANK-SET), consistent with prior studies (10) (**Figure 1A**). To determine if G9a and GLP form homo- or heterodimers, we co-expressed N-terminally 6xHis-tagged G9a (His:G9a) and N-terminally Maltose Binding Protein (MBP) tagged GLP or G9a (MBP:GLP or MBP:G9a) in *E. coli*. Dimerization is assessed by a sequential affinity purification strategy from *E. coli* lysates. Sequential cobalt and amylose resin purification of either the His:G9a::MBP:G9a homomeric or His:G9a::MBP:GLP heteromeric complexes retains a dimer (**Figure 1B**). Quantification of SyPRO Red stained bands indicated dimerization at 1:1 stoichiometry for both complexes (**Figure 1B**). Consistent with this observation, MBP:G9a or MBP:GLP constructs expressed individually had similar retention profiles through size exclusion chromatography (data not shown). We examined the stability of G9a homo- and G9a-GLP heterodimers using a dilution-based assay, which assesses the off-rate of the complex. Each complex was diluted to 40nM (about 100 times) and allowed to dissociate for 1-2 hours at room temperature. We assessed dimer association by precipitating the His-tagged protein with cobalt resin and determining the fraction of MBP protein that remained bound. We observed little dissociation in either G9a homo or heterodimers (**SFigure 1**). To exclude the possibility that the N-termini of G9a or GLP counteract complex formation, we examined complex retention following sequential affinity purification of full-length proteins produced in insect cells. We found them to behave similarly to ANK-SET (**SFigure 2A, B**). Together, these data indicate that G9a forms stable, 1:1 homodimers, as well as heterodimers with GLP.

**Figure 1:**
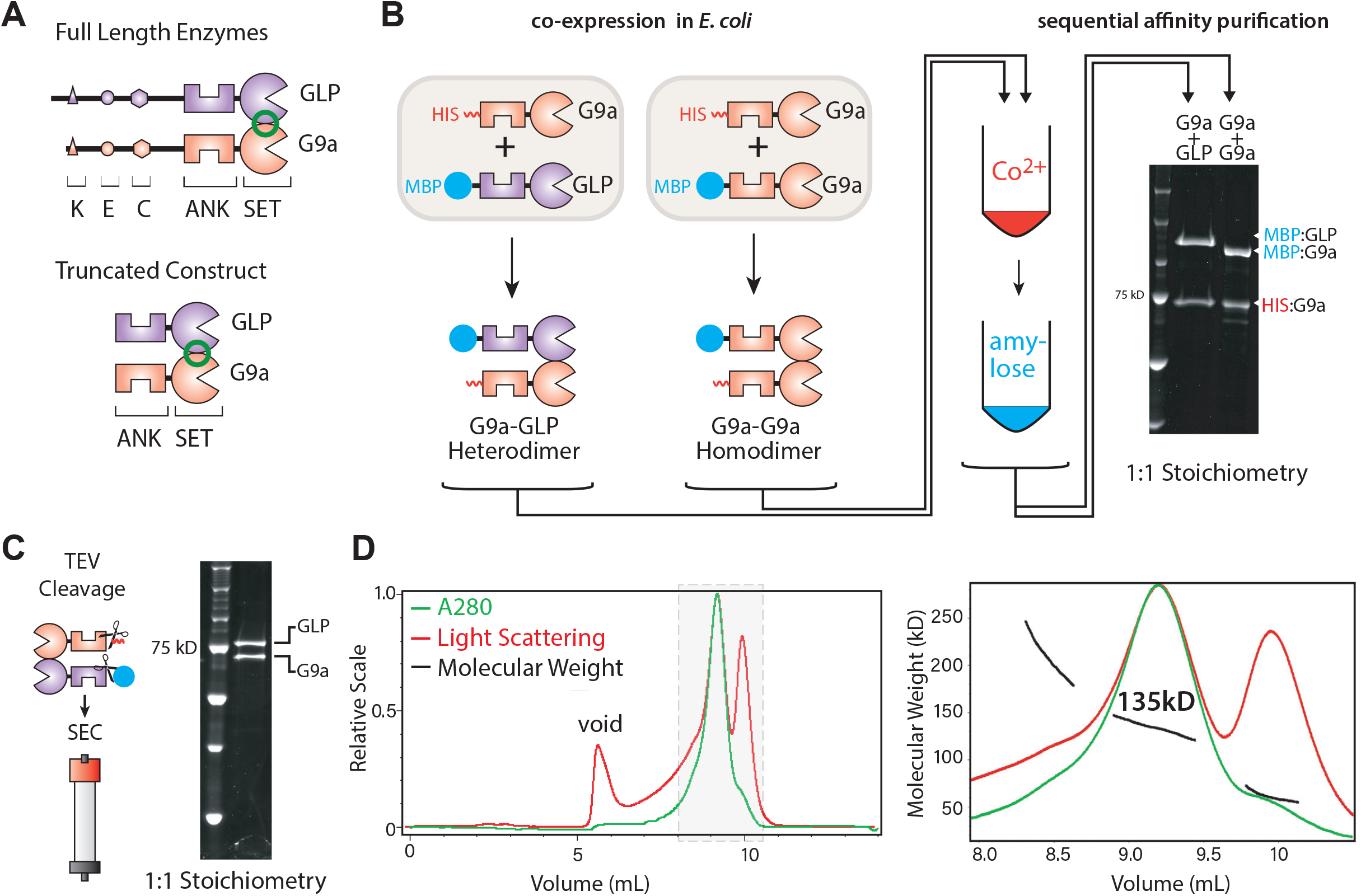
G9a and GLP form stable 1:1 homo- and heterodimers. **A**. Domain architecture of G9a-GLP. TOP: full-length enzymes. The N-terminus of G9a and GLP feature an automethylation residue (K), an acidic patch (E), and a cysteine-rich region (C). The C-termini contain ankyrin repeats (ANK) that bind H3K9me and a SET domain (SET) which is both the methylation catalytic domain and the dimerization interface (green circle). BOTTOM: Truncation ANK-SET construct used in this study. **B**. *E. coli* coexpression and purification strategy for identification of G9a homo and heterodimers. **C**. TEV cleavage of G9a-GLP heterodimers and purification via size exclusion chromatography **D**. SEC-MALS trace of G9a-GLP. LEFT: Full A280 (green) and Light Scattering (Red) traces. RIGHT: magnification of the main peaks with molecular weight determination (black). The measured molecular weight of the complex is 135kD (theoretical MW 142kD).

For further characterization of the G9a-GLP heterodimer, we removed both the His and MBP tags by TEV-mediated proteolysis (**Figure 1C**). Using this “tagless” G9a-GLP molecule, we next ascertained the number of enzymes per complex. To do so, we determined its molecular weight using size exclusion chromatography (SEC) followed by multi-angle light scattering (MALS). We observed one major peak in our SEC-MALS measurement (**Figure 1D** left) and determined it to have a molecular weight of ∼135 kDa **(Figure 1D**, right), roughly the theoretical molecular weight of our truncated G9a/GLP heterodimer (142 kDa). These data suggest that G9a-GLP is limited to a heterodimeric complex with one G9a and one GLP molecule.

### Heterodimerization facilitates G9a and GLP binding to H3K9 methyl peptides

The presumably monomeric ANK domains of G9a and GLP have been shown to interact with both H3K9me1 and H3K9me2 histone tails, to largely similar extents (7, 10). Using fluorescence polarization, we asked whether the ability to engage H3K9me1/me2 is preserved in the dimeric ANK-SET molecule and whether heterodimerization affected this reading function (**Figure 2**). Interestingly, except for GLP ANK-SET binding to me1 (**Figure 2A**), which is the tightest affinity we observe, the presence of the SET domain in the context of homodimers is broadly inhibitory to H3K9me binding compared to published ANK alone data (7, 10). GLP ANK-SET has 12.5-times reduced binding to me2 peptides (**Figure 2B**), and we confirmed that H3K9me binding interactions in the ANK-SET construct are dominated by the ANK domain (**Figure 2G,H**). Unlike GLP, G9a ANK-SET has no discernable affinity for me1 or me2 peptides (**Figure 2 C,D**), which we also observe for the full-length protein (**SFigure 2C**). Heterodimerization restored interaction with me1 and me2 to a range comparable to the ANK domain alone (**Figure 2 E,F**). Taken together, these results suggest that within the ANK-SET homodimeric context, the ANK domains of both G9a and GLP are partially compromised in their ability to bind methyl peptides and that this inhibition is overcome upon heterodimerization (**Figure 2I**).

**Figure 2:**
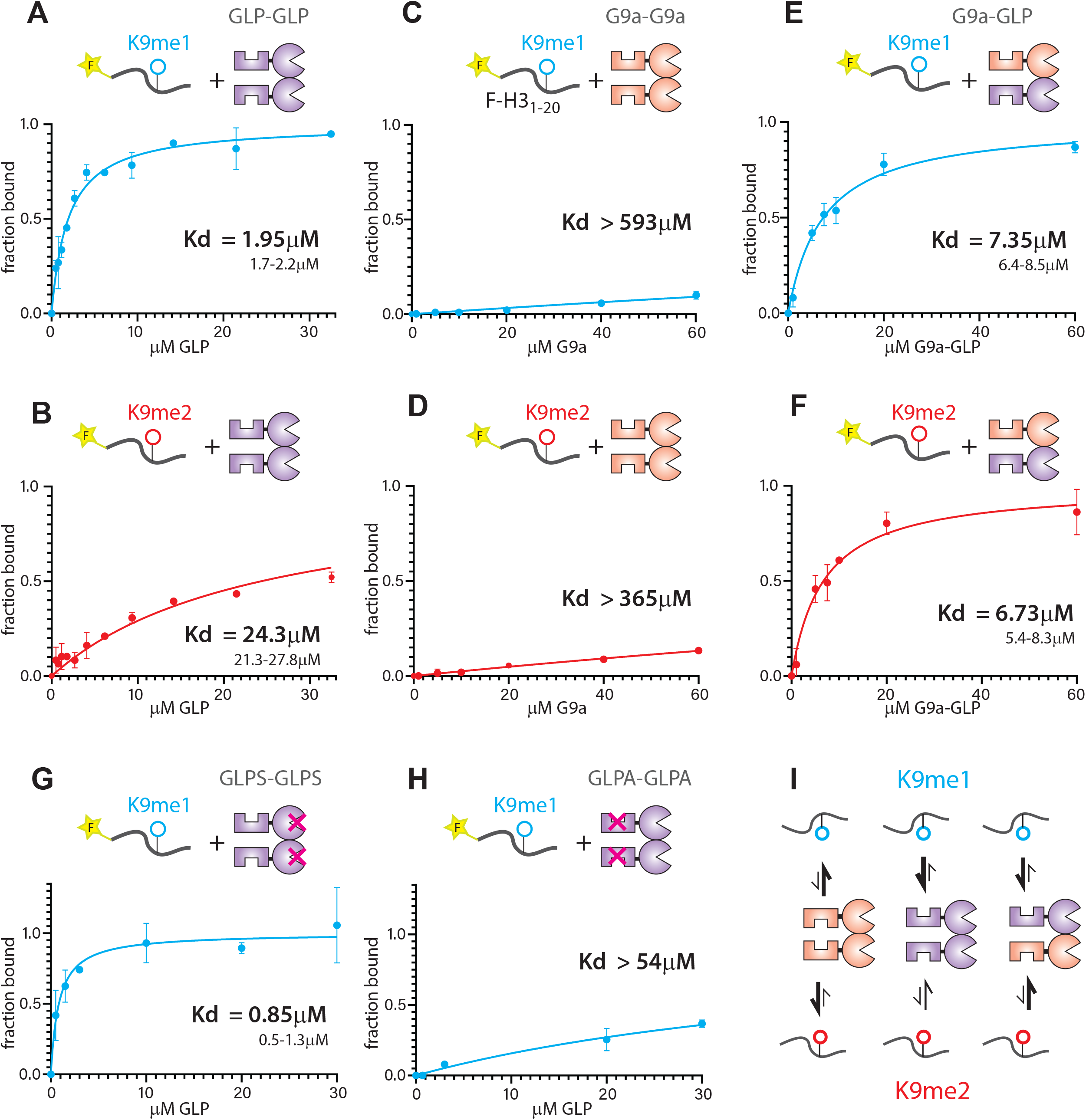
Heterodimerization facilitates G9a and GLP binding to H3K9 methyl peptides. Fluorescence Polarization measuring binding of H3K9 mono and dimethyl peptides to **A.&B**. GLP homodimers; **C.&D**. G9a homodimers; **E.&F**. G9a-GLP heterodimers; **G**. GLP homodimer with SET domain catalytic mutation (GLPS); **H**. GLP homodimer with ANK domain mutation (GLPA); K_d_ values are indicated on plots. The 95% confidence interval (CI) is shown for fits with significant binding saturation (A., E., F., G). For curves with limited saturation and fits with wide range of K_d_ values, the lower bound of the 95% CI is shown. **I**. Summary of peptide binding data. All error bars denote standard deviation from independent duplicate experiments.

### Heterodimerization Stimulates G9a and GLP Catalytic Activity

We next asked whether dimerization constitutively affects the methyltransferase activity of G9a-GLP. We determined Michaelis Menten kinetic parameters of G9a and GLP homodimers as well as the G9a-GLP heterodimer using an H3 histone tail peptide substrate (H3_1-20_, (11), **Figure 3A-C,J**)). While the K_M_ of all enzyme species were similar (**Figure 3J**) we observed changes of the k_cat_ within the heterodimer beyond those expected from equal contributions from G9a and GLP: The G9a and GLP k_cats_ on H3_1-20_ are ∼33min^-1^ and ∼14min^-1^, respectively. Rather than an intermediate value, G9a-GLP’s k_cat_ is ∼33min^-1^, suggesting one or both enzymes are more active in the heterodimer than their respective homodimers. To determine which enzyme is stimulated in the heterodimer we made point mutations in either G9a (G9aS, Y1120V, Y1207F) or GLP (GLPS, Y1240F) abrogating their catalytic activity while still allowing dimerization with a wild type allele of their binding partner (15, 18). Comparing G9aS-GLP to GLP-GLP we observe that GLP’s k_cat_ increased ∼2 times upon heterodimerization (**Figure 3C**). Activity comparisons of G9a-GLPSand G9a-G9a suggested very minor increase in activity of G9a in the heterodimer even at saturating peptide concentrations (**SFigure 4C**), indicating that only GLP’s activity is meaningfully enhanced in the heterodimer. Thus, these data imply a modest catalytic enhancement in G9a-GLP versus the homodimeric forms.

**Figure 3:**
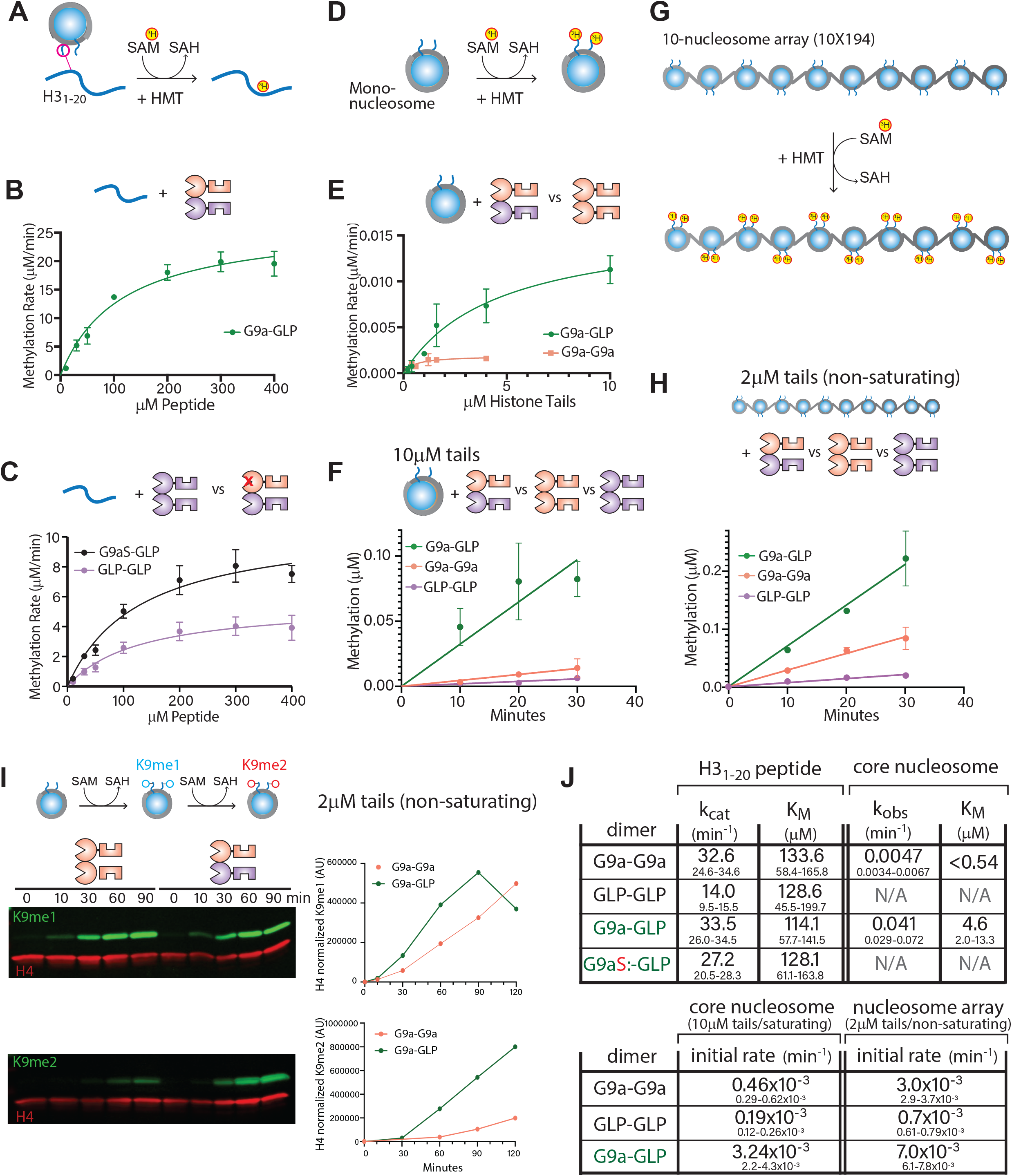
Heterodimerization stimulates G9a and GLP catalytic activity. **A**. Reaction scheme of H3 tail peptide methylation. SAM cofactor was labeled with tritium (^3^H). **B**. Michaelis Menten fit of G9a-GLP heterodimer methylating H3 tail peptide. **C**. Michaelis Menten fit of GLP homodimer (purple) vs G9aS-GLP heterodimer (black). **D**. Reaction scheme of mononucleosome methylation reactions. **E**. Michaelis Menten fit of G9a-GLP heterodimer (green) and G9a homodimer (pink) methylating mononucleosomes. **F**. Initial rate comparison of G9a-GLP (green), G9a homodimer (pink), and GLP homodimer (purple) methylating mononucleosomes under saturating conditions (10μM histone tails; 5μM mononucleosome). **G**. Reaction scheme of nucleosome array methylation reactions. **H**. Initial rate comparison of G9a-GLP (green), G9a homodimer (pink), and GLP homodimer (purple) methylating 10-nucleosome arrays under non-saturating conditions (2μM histone tails; 1μM nucleosomes). **I**. LEFT: Western blot time course of H3K9me1 or H3K9me2 (green) production on mononucleosomes under non-saturating conditions (2μM histone tails; 1μM mononucleosomes). H4 was blotted for as an internal control (red). RIGHT: The H4-normalized production of H3K9me1 or me2 by G9a (pink) or G9a-GLP (green) is plotted over time. **J**. Kinetic parameters for histone peptide and nucleosome methylation. Values reflect kinetic parameters (upper value) and 95% confidence intervals (lower values). k_obs_ for nucleosomes denotes a first order rate constant under conditions where the SAM pocket is not saturated (see SFigure 3A,B). All error bars denote standard deviation from independent duplicate experiments.

Next, we asked if the G9a-GLP heterodimer exhibits altered methylation kinetics on a mononucleosome, which mimics its cellular chromatin substrate (**Figure 3D**). We first measured Michaelis Menten parameters under multiple turnover conditions, just like for H3_1-20_, for G9a-GLP and G9a-G9a on a reconstituted mononucleosome. In striking contrast to our observations on H3_1-20_, G9a-GLP’s k_obs_ is 10 times higher than that of G9a-G9a (**Figure 3E,J** and **SFigure 4B**), which we confirmed measuring initial rates under near-saturating conditions for mononucleosomes. Under these conditions, the heterodimer also exhibits higher activity (∼ 8 times higher k_obs_) than G9a-G9a, with the GLP homodimer the slowest of the three enzyme constructs (**Figure 3F**). We repeated this initial rate comparison for 10-nucleosome arrays (**Figure 3G**), to test whether this enhancement holds for chromatin substrates. However, due to technical limitations we could do so only under non-saturating conditions for G9a-GLP (2mM histone tails). Nevertheless, even in this condition G9a-GLP was the fastest of the three enzymes, and GLP the slowest, as for the mononucleosome case (**Figure 3H**). Surprisingly, while G9a-GLP accelerates H3K9me1 and me2 production on nucleosomes versus G9a, the overall rate enhancement appears to be coded most strongly in the H3K9me1-H3K9me2 transition (**Figure 3I**). We conclude that chromatin substrates bring to fore the intrinsic catalytic differences between homo and heterodimers, revealing a dramatically increased k_obs_ for G9a-GLP. We note that, concomitant with this k_obs_ increase, however, we also noticed an increase in the nucleosome K_M_ for G9a-GLP (**Figure 3E,J**). Joint increases or decreases in k_cat_ and K_M_ on chromatin substrates have been observed before and can be indicative of changes in the number of substate encounter modes compared to simpler substrates (11).

### The Heterodimer interacts with chromatin in a G9a-specific manner, but unlike G9a, recognizes methylated nucleosomes

As in the studies above with the H3 tail peptide, we used fluorescence polarization to measure the binding affinity of each ANK-SET dimer on unmethylated, or H3K9 mono-or dimethylated mononucleosomes. On the unmethylated nucleosome, for G9a-G9a and G9a-GLP homo- and heterodimers, respectively, we measure a similar K_d_ in the range of ∼6-7µM (**Figure 4 A, C**). For G9a, this interaction requires a catalytically active SET domain (**SFigure 5A**). GLP-GLP exhibits a significantly lower affinity, with a K_d_ of 26.3µM (**Figure 4B**), indicating that binding to the unmethylated nucleosome in the G9a-GLP heterodimer is mostly driven by G9a. Binding of unmethylated nucleosomes by G9a-GLP does not appear to be depend on contacts with DNA, unlike for other HMTs (11, 12), as G9a-GLP does not show appreciable binding to DNA compared to Clr4 HMT control (**SFigure 6**). However, DNA binding may be conferred by regions of G9a or GLP outside the ANK-SET module, in the protein N-terminus (**Figure 1A**). Thus, the engagement mode of G9a and G9a-GLP with unmethylated nucleosomes seems to require interaction with the H3 tail via the SET domain active site. It is possible that the loss of engagement with the nucleosome in G9aS Y1120V/Y1207F is due to more extensive rearrangements in the G9a SET domain H3 binding pocket mutants. Even though the data points to the SET domain-H3 tail contact as necessary for nucleosome engagement, given the increased affinity for nucleosome over H3 peptides (see **SFigure 2C, D**), G9a/ G9a-GLP likely make additional contacts to the histone octamer.

**Figure 4:**
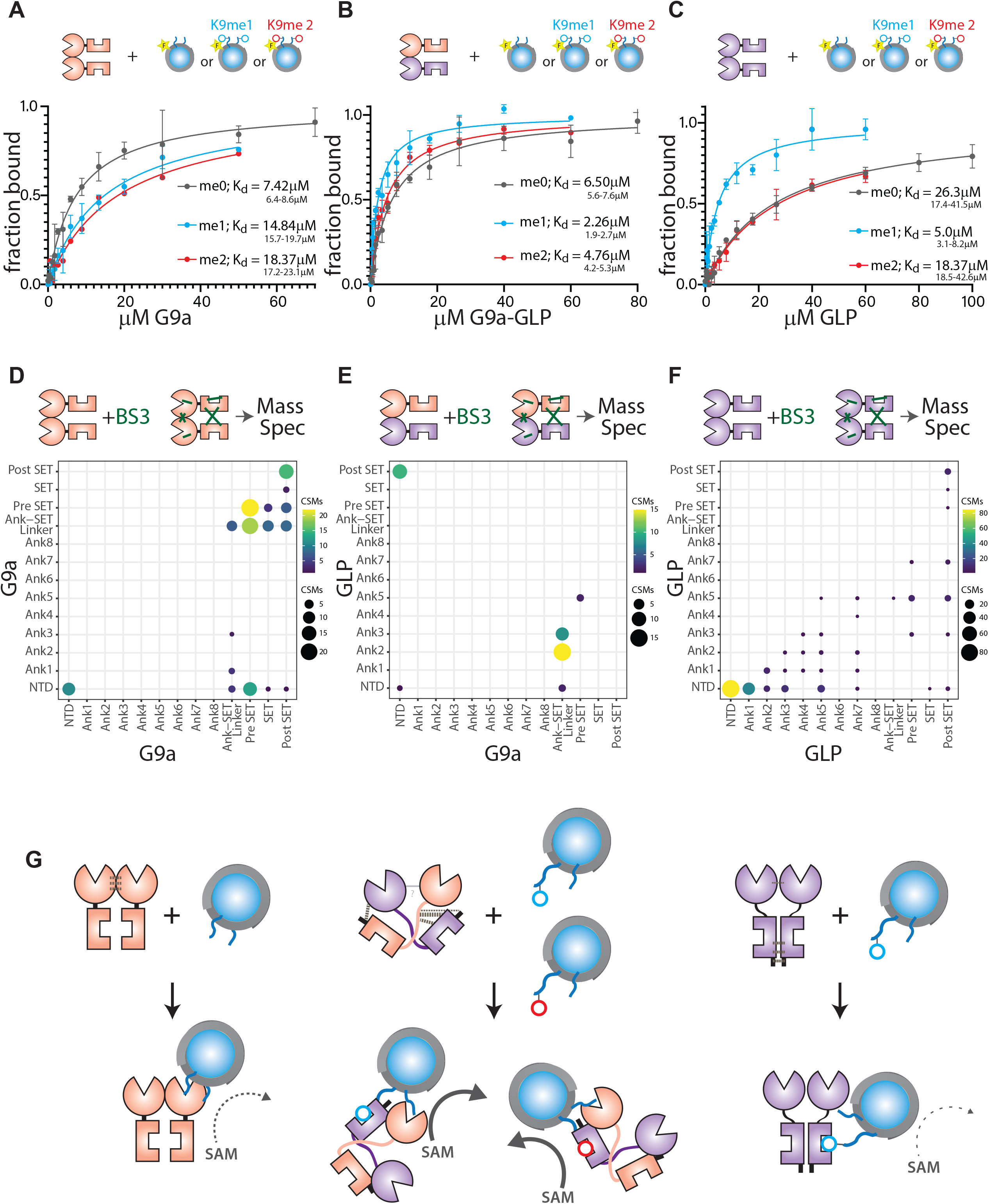
Unique nucleosomes engagement and ground statement conformation of G9a-GLP heterodimer. **A.-C**. Fluorescence polarization of unmethylated (H3K9me0; grey), H3K9me1 monomethylated (teal), and H3K9me2 dimethyated (red) mononucleosomes containing 147bp ‘601’ DNA 5′ fluorescein-labeled bound to **A**. G9a **B**. G9a-GLP **C**. GLP-GLP. The 95% confidence interval (CI) is shown for all K_d_s. **D.-F**. Crosslinking Mass Spectrometry for G9a (**D**), G9a-GLP (**E**), and GLP (**F**). Dot size and color indicates number of crosslinked spectral matches (CSM) summarized for domain pairs. Note that G9a and GLP data by necessity include intra-and inter protomer crosslinks. **G**. Model. G9a protomers interact predominantly via the SET domain. The ANK domains are in an auto-inhibited state. Nucleosome engagement is via the H3 tail. G9a-GLP makes cross-protomer N- to C-terminal contacts, leading to a different ground-state conformation with the ANK domains free. Either G9a and GLP (shown) ANK could engage with H3K9me1 or me2 nucleosomes. The engagement mode of G9a-GLP with the nucleosome results in increased catalysis. GLP associates strongly at the N-terminus, allowing the ANK domains to access methylated residues. However, GLP has limited engagement with the nucleosome via the SET domain, resulting in slow catalysis.

We also assessed how H3K9 mono, and di-methylation impacts the association of G9a, GLP or G9a-GLP to nucleosomes. Consistent with G9a’s inability to bind H3K9me1 or me2 tail peptides, the affinity of G9a to H3K9me1 and me2 nucleosomes decreases compared to unmethylated nucleosomes (**Figure 4A**). This would also be consistent with the primary nucleosome interaction mode requiring the SET domain catalytic domain, which binds methylated tails much more weakly than the ANK domain (**Figure 2G, H**). GLP is able to bind relatively tightly to H3K9me1, but not H3K9me2 nucleosomes (**Figure 4B**). Since GLP binds H3K9me1 peptides tightly, this may suggest that GLP can engage with the nucleosome via ANK domains when H3K9 is monomethylated, but only weakly via the SET domain in the unmethylated state. In contrast, the G9a-GLP heterodimer exhibits mild enhancement of binding H3K9me1 and me2 nucleosomes over the unmethylated state. However, since fluorescence polarization cannot distinguish the engagement mode of G9a-GLP, we cannot conclude to what degree ANK and SET engagement contributes to these overall K_d_s. Nevertheless, only G9a-GLP can apparently recognizes both H3K9me1 and H3K9me2 on nucleosomes.

### G9a, GLP and G9a-GLP exhibit unique inter and intra molecular contacts

The unique catalytic and substrate-binding behaviors of G9a-GLP, in particular on chromatin, may be encoded by a different ground state conformation for the heterodimer, or by induced changes that occur upon substate engagement and that are not evident in the ground state. To distinguish between these possibilities, we conducted BS3 crosslinking followed by Mass Spectrometry of G9a and GLP homodimers and G9a-GLP heterodimer. We note that we may not be able to detect all protein-protein contacts via this method (see Supporting discussion). Nonetheless, we observed marked differences between all three dimers. Crosslinks in G9a are clustered in the Set domain, most prominently in the “pre-SET” and “post-SET” region (**Figure 4D**). While GLP also displayed contacts in the SET domain, it displayed much more prominent crosslinks at the N-terminus of the ANK-SET construct, upstream of the ANK (NTD, **Figure 4F**). We note that for both G9a and GLP, we cannot differentiate whether these contacts are intra- or intersubunit. However, we can infer that the most frequent contacts are largely within the same region of the protein, N terminus/ANK versus C terminus (ANK-SET linker and pre-SET, SET and post-SET). For G9a-GLP, we can specifically isolate inter-subunit crosslinks, as we can filter out intra-subunits crosslinks, which results in overall isolating fewer crosslinks compared to the homomeric dimers. Surprisingly, we find a different pattern of crosslinks for G9a-GLP, where the strongest crosslinks are from the N-terminus of G9a to the post-SET of GLP, or from the ANK of GLP to the ANK-SET linker or pre-SET of G9a. This is qualitatively different from G9a or GLP, pointing to a different ground state conformation. A repeat of this experiment with the MBP-tagged proteins yielded a similar overall picture (**SFigure 7**), though in this case, we uncovered serval MBP crosslinks which may have biased the overall distribution of captured contacts.

## Discussion

Unusual among metazoan histone methyltransferases, G9a and GLP form dimers *in vivo* and this dimerization is essential for function (9). We discuss our findings regarding the ability of these proteins to 1. homo-and heterodimerize and the unanticipated consequences of heterodimerization on both H3K9me 2. reading, and 3. writing.

1. *Constitutive dimerization:* The degree to which dimerization, or the stability of dimers, is regulated by homo-and heteromeric associations is unknown. Our data argue that the ANK-SET portion of G9a and GLP homo- and heteromeric species form 1:1 dimers with low apparent dissociation and that efficient dimerization is not limited in the homomeric context (**Figure 1, SFigure 1**). However, *in vivo* experiments indicate that when G9a and GLP are present at equal cellular stoichiometry, the heterodimer is the preferred form and that homodimers can form when G9a or GLP is in excess (9). This *in vivo* preference may be due to regulation outside the ANK-SET domain. Taken together with our results, a picture emerges where G9a and GLP constitutively form homo- or heterodimers with relative pools determined by the steady-state accumulation of either protein. Homodimers, when formed, may then execute unique functions, for example in DNA repair for GLP (19), and in terminal muscle differentiation, where G9a and GLP control non-overlapping genes sets (20). We speculate that the heterodimer, with increased catalysis on chromatin and product recognition (below), is required when large domains of H3K9me2 are first formed and then maintained through cell division (5, 6), and that specialized and local chromatin methylation, or methylation of non-histone targets, may be carried out by homodimers.
2. *De-inhibition of H3K9me2 reading:* Our data suggest that H3K9me reading, in particular H3K9me2 is inhibited within the ANK-SET homodimer. Only the heterodimer appears capable of strong H3K9me2 recognition (**Figure 2**), while both GLP-GLP and G9a-GLP can recognize H3K9me1. This differs from findings on the presumably monomeric G9a and GLP ANK domains alone as measured by fluorescence polarization (7) or ITC (10). Because the affinities we measure for H3K9me1/2 binding, where observed, are within 2-fold of these published affinities for ANK alone (7, 10), we do not believe that the ANK domains in our hands have lower specific activity. Instead, we interpret our data to indicate that either 1. inhibitory contacts between ANK and SET, or ANK and ANK across the two dimers are alleviated in G9a-GLP or 2. Heterodimerization induces a conformational change within the ANK. Our crosslinking-Mass spectrometry results favor differences in the ground state conformation of the heterodimer, which may “liberate” the ANK domains of G9a and possibly GLP to engage with H3K9me (**Figure 4D-F**, model **Figure 4G**). Beyond these interactions in the ground state where the protein has not engaged substrate, H3K9me ANK binding sites may be initially unavailable in the naïve protein but become fully available following initial nucleosome engagement to bind adjacent H3K9me sites. This could account for the observation of H3K9me1 or me2 specific stimulation of methylation by G9a and GLP homodimers on chromatin circles (10).
3. *Stimulation of chromatin methylation:* We observe both modest, constitutive activation of GLP on H3 tail peptides, and a dramatic increase in the k_obs_ towards nucleosomes compared to the homodimers. We believe two mechanisms can account for these results:
  a. Relief of general autoinhibition. Like other histone methyltransferases, G9a and GLP may be limited by autoinhibition. The fission yeast Clr4 enzyme contains an autoinhibitory loop that is relieved by automethylation (21) and similar inhibitory loops have been shown for PRC2 and NSD1 (22, 23). The constitutive activation we observed in GLP upon heterodimerization (**Figure 3**) may be due to the restructuring of an autoinhibitory loop in the post-SET domain (24). Additionally, the inability of G9a to recognize H3K9 methylation on peptides or nucleosomes (**Figures 2&4**) suggest that the ANK of G9a is in an inaccessible state.
  b. Optimized modes of substate engagement. The observation that GLP binds poorly to nucleosomes indicates that G9a and G9a-GLP feature more optimal nucleosome engagement specifically, although this is not the case for the peptide, as the K_M_s of all three enzymes are similar. Therefore, while engagement of the SET domain with the H3 peptide is critical for nucleosome binding (**SFigure 5A**), additional contacts are likely made by G9a/G9a-GLP to other surfaces of the histone octamer. Further, G9a-GLP likely features a different ground state conformation that may enable more productive engagement with the nucleosome compared to G9a, resulting in an increased k_obs_. Given that G9a’s K_M_ is significantly lower than its nucleosomal K_d_ (unlike in the G9a-GLP case), this inefficient nucleosome catalysis could be explained by either an increase in non-productive binding modes (11), or inefficient product release (25). While we cannot determine the true k_cat_ for nucleosomes given signal limitation (see methods) we note that the specificity constant (k_cat_/K_M_) is not raised compared to peptides, at least on mononucleosomes. However, the elevated k_obs_ observed on nucleosome arrays compared to mononucleosomes, even at sub-saturating conditions (**Figure 3J**), hints that G9a-GLP’s true k_cat_ on chromatin may still be higher.

The unique ability of the G9a-GLP molecule to recognize H3K9me2 as well as methylate chromatin, especially compared to the G9a homodimer (**Figure 4G**), begins to explain the strong requirement of the heterodimer for global maintenance of H3K9me2 and normal embryogenesis.

## Methods

### Purification of G9a/GLP Homo and Heterodimers

To isolate the ANK-SET (10) G9a-GLP heterodimer, we coexpressed N-terminally tagged His:G9a and MBP:GLP from a single plasmid (QB3 Berkeley Macrolab expression vectors) in *E. coli* DE3 Rosetta cells and performed sequential cobalt- and amylose-charged resin affinity chromatography purification. Cells co-expressing His and MBP constructs were lysed on ice via sonication in lysis buffer (100mM Tris pH 8, 300mM NaCl, 10% glycerol (v/v), 0.1% Tween-20, with freshly added 1mM β-mercaptoethanol (BME), 1mM PMSF, 5mM Benzamidine, 200µM Leupeptin, Aproptinin, Pepstatin, Phenantroline). The clarified lysate was then bound to cobalt-charged resin (Takara) for 1hr and washed twice with lysis buffer. His tagged proteins were eluted with lysis buffer containing 400mM imidazole and bound immediately to amylose resin (NEB) for 1hr. MBP tagged proteins were eluted with lysis buffer + 20mM maltose. Affinity tags were then removed by incubation with 12mg TEV protease for 1hr at 25°C. TEV protease was absorbed to cobalt resin and the cleaved heterodimer was further purified by size exclusion chromatography (Superdex 200 Increase 10/300 column) and buffer exchanged into storage buffer (100mM Tris pH 8, 100mM KCl, 10% glycerol, 1mM MgCl_2_, 20µM ZnSO_4_, 10mM BME). Homodimeric MBP:G9a or MBP:GLP ANK-SET constructs were purified as above, omitting the cobalt resin purification. All protein constructs were quantified using SDS page with BSA standards and Sypro Red stain.

### Size-Exclusion Chromatography coupled to Multi Angle Light Scattering

For size-exclusion, protein samples were injected at 10µM into a silica gel KW804 chromatography column (Shodex; Shanghai, China). For MALS, the chromatography system was coupled to an 18-angle light-scattering detector (DAWN HELEOS II, Wyatt Technology; Santa Barbara, CA) and a differential refractometer (Optilab-rEX, Wyatt Technology). Data collection was done at a flow rate of 0.4mL per minute. SEC MALS data were collected and analyzed using Astra 7 software (Wyatt technology; Santa Barbara, CA).

### Preparation of mononucleosome substrates

Histone proteins and nucleosomes were purified as described (26) with the following modifications: Following assembly nucleosomes were dialyzed overnight into storage buffer (above) or FP storage buffer (50mM HEPES pH 7.5, 100mM KCl, 10% glycerol) depending on the application. Dialyzed samples were concentrated using 10kda Millipore Sigma Amicon Ultra Centrifugal Filter Units. For fluorescence polarization nucleosomes were reconstituted with a 5′ fluoresceinated DNA template. H3K9 dimethylation was installed via Methyl-Lysine Analog (MLA) technology as described (10, 26). K9 MLA-monomethylated H3 was purchased from Active Motif. H3K27A was introduced by site directed mutagenesis and K27A mononucleosomes produced as above.

### Methylation Assays

All methylation reactions used a tritium-based assay to detect the methylated product. Substrate peptides or nucleosomes were mixed in a solution containing 9µM tritiated S-Adenosyl Methionine (SAM, Perkin Elmer) cofactor, and reactions were initiated upon addition of 0.4-0.8µM enzyme. Before adding enzyme, peptide reactions were supplemented with 1000□M cold SAM to fully saturate the SAM binding pocket (S Figure 3). Due to signal limitations, nucleosome reactions were not supplemented with SAM (see below). Reactions were run in 100mM Tris pH 8, 100mM KCl, 1mM MgCl_2_, 20µM ZnSO_4_, 10mM BME and quenched with laemmli buffer. Peptide reactions were performed as described (11). Mononucleosome methylation reactions were read out via autoradiography. Proteins were separated on a 18% SDS PAGE gel which was dried and exposed to a GE Tritium Phosphor screen for 72 hours along with a standard curve of tritiated SAM spotted on Whatman paper, and imaged on a STORM imager. Images were quantified using Image Quant software (Cytiva).

### Kinetics

All methylation reactions (except SFigure 4A)were performed under Multiple Turnover conditions (25). G9a and GLP each catalyze a two-substrate bi-bi reaction using the cofactor SAM and H3K9 as substrates (16). To determine initial rates, methylation time courses were traced at various concentrations of H3K9-containing substrate while keeping the concentration of SAM and enzyme constant. Plots of initial rate vs. concentration H3K9 substrate measured in duplicate or triplicate were fit to V = Vmax*[S]/(K_M_ + [S]) using Prism software to extract K_M_ and k_cat_ (or k_obs_) values as well as the 95% confidence interval bounds of the fit. For peptide reactions, we saturated the SAM binding pocket (see above) with a mixture of tritiated and non-tritiated SAM to measure kinetic parameters under pseudo first-order conditions. For nucleosome experiments, we could not fully saturate the SAM pocket, due to signal limitations, hence we report k_obs_ values for nucleosomes.

### Fluorescence polarization

Polarization assays with fluoresceinated H3_1-20_ K9me1 or K9me2 peptides, 147bp 601 DNA template, and core nucleosomes were performed and K_d_ values estimated as described (11) with the following modifications: The reaction buffer was 100mM Tris pH 8, 100mM KCl, 1mM MgCl_2_, 20µM ZnSO_4_, 10mM BME, 0.1% NP-40. The concentration of peptides and nucleosomes was 150nM and 200nM for DNA. Polarization measurements were conducted on a Biotek Cytation 5 (peptide and nucleosome) or Molecular Devices Spectramax (DNA) plate reader with low volume plates (Corning) in either 10µl (peptide, nucleosome) or 40µl (DNA) total volume.

### Crosslinking Mass Spectrometry (CLMS)

GLP, G9a or G9a-GLP were either cleaved from His/MBP tags (Figure 4) or left uncleaved (SFigure 7) and dialyzed into crosslinking buffer (25mM HEPES pH 7.5, 140mM KCl, 0.5mM MgCl_2_, 1mM BME). CLMS was generally performed as described (27) with minor modifications: Crosslinking was performed at 4mM monomer and 0.75mM BS3 crosslinker for 33min at room temperature. The reaction was quenched with Tris HCl pH 7.5. Crosslinking was confirmed via SDS-PAGE. Total protein was precipitated with cold acetone. Size exclusion fractions enriched in crosslinked peptides were combined into two mass spectrometry samples and each one was acquired over a 90 minute UPLC gradient acquiring sequential HCD and EThcD product ion spectra on a Fusion Lumos mass spectrometer (Thermo Scientific). Peaklists were generated using PAVA for each dissociation method and searched with Protein Prospector 6.3.23. The search database included sequences for human G9a and GLP methyltransferases, as well as four contaminating proteins detected in moderate abundance (from *e coli* chaperones and bovine protein standards). A randomized fasta database that was 10x longer than the target database was used for FDR estimation. Only sample specific protein crosslinks are reported. Crosslinked spectral matches (CSMs) were classified by picking an SVM.score threshold corresponding to at a 1% false discovery rate (FDR). CSMs are reported at the unique-residue-pair level in Supplemental Table 1 and plotted at the domain-pair level in Figure 4 and SFigure 7. Domain pair reporting was done by counting the number of product ion spectra above the SVM.score threshold (eg CSMs) that mapped to each domain pair. CLMS classification and reporting was performed using Touchstone, an in-house R library.

## Supporting information

Source data for Crosslinking Mass Spectrometry data in Figure 4

## Acknowledgments

We thank Carol A Gross and Danica Fujimori for thoughtful discussions and feedback. We thank the Southworth lab for training and access to the SEC-MALS equipment. N.A.S. was supported by a Hooper Graduate Student Fellowship, B.A-S. was supported by grants from the National Institutes of Health (DP2GM123484 and R35GM141888) and the University of California, San Francisco Program for Breakthrough Biomedical Research. Mass spectrometry experiments were supported by the Adelson Medical Research Foundation and the University of California, San Francisco Program for Breakthrough Biomedical Research.

## Competing interests

The authors declare no competing interests.

## Supporting material

### Supporting Methods

#### Dilution Experiment

Purified His:G9a::MBP:GLP or His:G9a::MBP:G9a complexes were diluted to 40nM in lysis buffer and allowed to dissociate for 1-2hr at room temperature in a volume of 20µL. 7µL cobalt-charged resin was added to each sample and incubated for 30min to allow resin binding. The resin was washed twice with lysis buffer and 20µL 1x Laemmli Buffer was then added to resin. The ratio of MBP tagged protein to his tagged protein was assessed via SDS page with BSA standards and Sypro Red stain. Fraction assembled was =1 for a “stock protein” (SP) control that was not put through this assay and had a concentration of >4µM.

#### Insect Cell Expression

Bacculovirus containing full-length His:G9a and STREP:GLP co-expression cassettes (QB3 Berkeley Macrolab) were used to infect Sf9 cells (Expression Systems, Davis, California) grown in ESF921 media. Cells were infected for 72hr at an MOI of 0.1. Infected cells were flash frozen and stored at -80°C until thawed for purification. Cells were lysed on ice via sonication in lysis buffer (100mM Tris pH 8, 300mM NaCl, 10% glycerol (v/v), 0.1% Tween-20, with freshly added 1mM β-mercaptoethanol (BME), 1mM PMSF, 5mM Benzamidine, 200µM Leupeptin, Aproptinin, Pepstatin, Phenantroline). The clarified lysate was then bound to streptactin superflow resin (IBA) for 1hr and washed twice with 100mM Tris pH 8, 750mM NaCl, 10% glycerol (v/v), 0.1% Tween-20, freshly added 1mM BME. Strep tagged proteins were eluted with lysis buffer containing 5mM desthiobiotin and bound immediately to cobalt-charged resin (Takara) for 1hr. His tagged proteins were eluted with lysis buffer + 400mM imidazole. Complex formation was assessed via western blot.

#### Electromobility Shift Assay

To assess binding to 147bp ‘601’ DNA via EMSA, G9a-GLP or Clr4 were incubated with 200nM of a 147bp 601 DNA template in 50mM HEPES pH 7.5, 100mM KCl, 10% glycerol, 0.1% NP-40 in a 10µL volume. Upon mixing, 50% glycerol was added to 5% final and samples were run on a 5% Tris-Glycine Native Gel. Samples were imaged using Chemidoc MP Imaging System (Biorad).

#### Expression in E. coli and purification of full-length G9a and SET domain G9a-GLP heterodimer

The purification strategy for SET domain G9a-GLP heterodimer was as for the ANK-SET version, with identical purification affinity tags. Full-length human G9a was cloned with a N-terminal MBP tag (MBP:FL-G9a), analogous to ANK-SET MBP:GLP and purified as above, without cleavage of the MBP tag to preserve maximal yield.

### Supporting discussion on Crosslinking Mass Spectrometry of G9a, GLP, and G9a-GLP

In this experiment, we crosslinked all three dimers with the BS3 crosslinker. One surprising finding for G9a-GLP is the absence of specific G9a-GLP crosslinks via the SET domain. Mutations in the SET domain have been shown to disrupt heterodimers *in vivo* (15) and in HEK cells, SET domains of G9a and GLP have been shown to co-immunoprecipitate (9). It is possible we did not observe crosslinks at these sites for the following technical reasons: 1. For G9a-GLP, crosslinks are excluded that occur over identical sequences in both proteins, as those cannot be assigned. For example, G9a and GLP share the crosslinkable sequence CWYDKDGR in the pre-SET domain. Other short identical sequences exist that would not be assigned should those result in crosslinks. The CLMS scoring algorithm prioritizes intra-protein results over analogous inter-protein results so CSMs matching these peptide would be reported as either G9a-G9a or GLP-GLP crosslinks. 2. It is possible that our crosslinking was not complete. We chose a concentration of BS3 crosslinker (0.75mM) that results trapping of a dimeric form and some minor accumulation higher molecular weight forms, but not significant accumulation of aggregates. Under these conditions, the monomeric form is not fully depleted. Hence it is possible that potential contacts did not get trapped by crosslinking at this BS3 concentration. 3. It is possible we did not identify a particular crosslink because the physiochemical properties of the crosslinked peptides result in poor ionization or chromatography such that the precursor ion is not selected for MS2 in a complex mixture.

**Supporting Figure 1:**
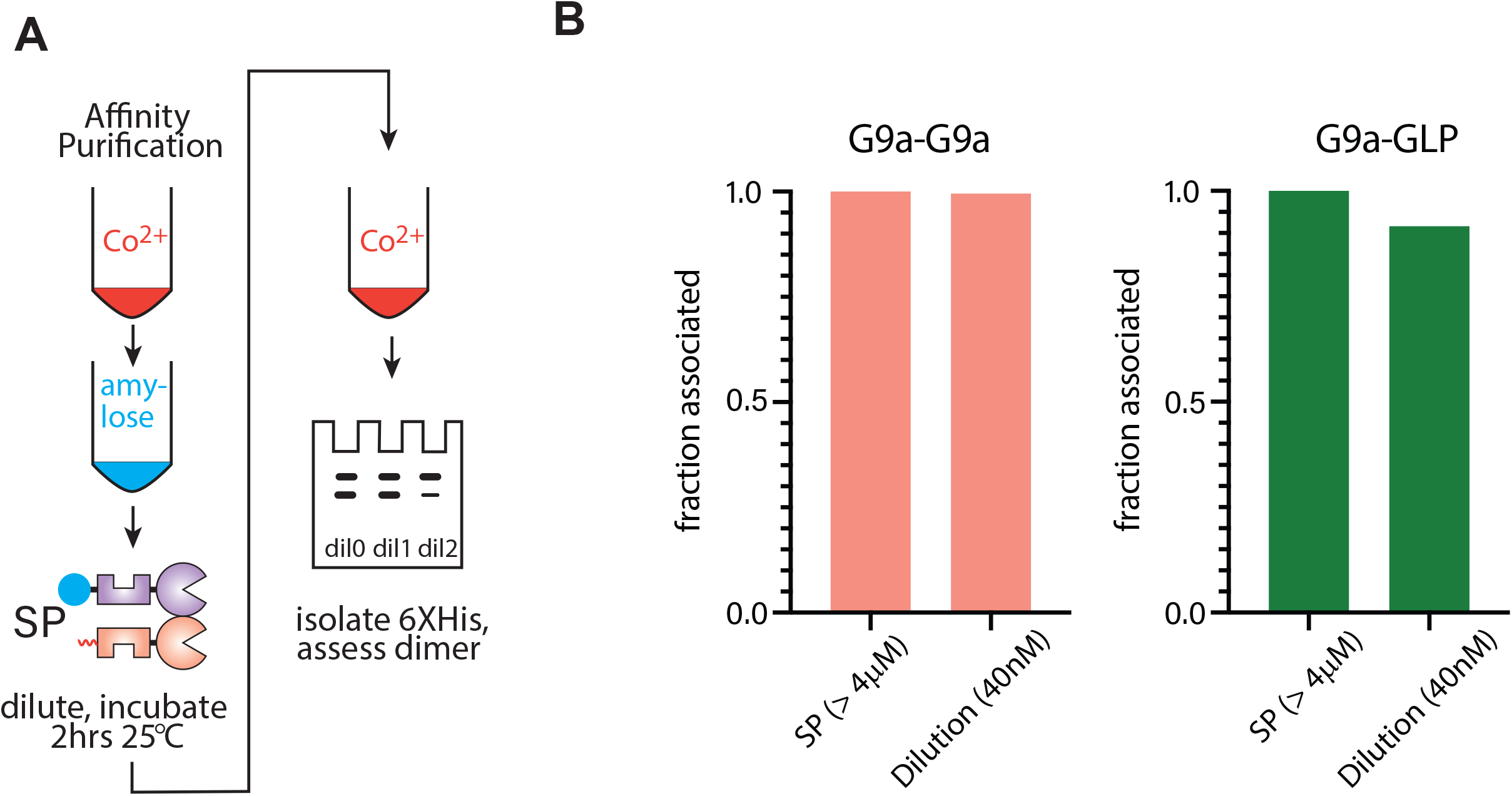
Stability of G9a-G9a and G9a-GLP complexes through a dilution assay. **A**. Experiment scheme. His:G9a coexpressed with either MBP:G9a or MBP:GLP was purified by sequential affinity purification as in Figure 1. The complexes were diluted and kept at 25°C for 1-2hrs. After this incubation, His:G9a was isolated via cobalt resin precipitation. **B**. The relative amount of His and MBP tagged proteins retained after cobalt resin precipitation was quantified by SyPRO Red staining and normalized to a stock protein (SP, > 4μM) of purified undiluted protein (fraction associated). The highest dilution for G9a-G9a and G9a-GLP, 40nM or > 100 times dilution, is shown.

**Supporting Figure 2:**
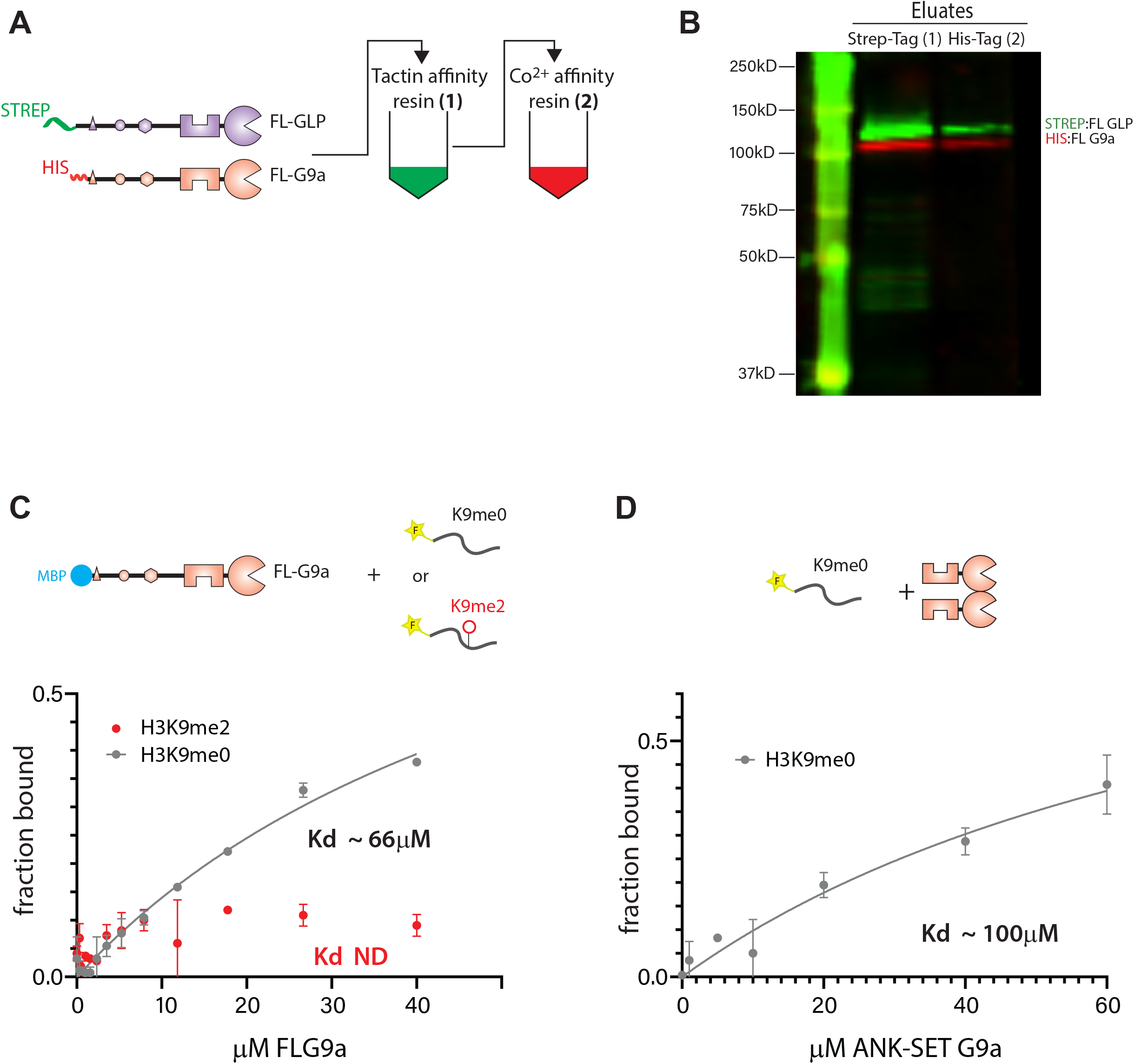
Behavior of Full-length G9a and GLP. **A**. Full-length GLP and G9a were N-terminally tagged with a STREP or 6XHis tag, respectively, co-expressed from a baculoviral construct in Sf9 insect cells, and isolated by sequential STREPTACTIN (Tactin) and Cobalt (Co^2+^) affinity resins. **B**. Western blots of baculovirus-infected or uninfected Sf9 lysates (LEFT) or affinity resin eluates (RIGHT) with anti-His tag or anti-STREP tag antisera. The single-channel and merged images are shown. **C**. Fluorescence polarization with *E. coli* expressed MBP:FL-G9a. The estimated (due to lack of saturation) K_d_ for H3K9me0 is shown, the K_d_ for H3K9me2 could not be determined. **D**. Fluorescence polarization with ANK-SET G9a and H3K9me0. The estimated (due to lack of saturation) K_d_ is shown. Error bars in C. and D. indicate standard deviation of two repeats.

**Supporting Figure 3:**
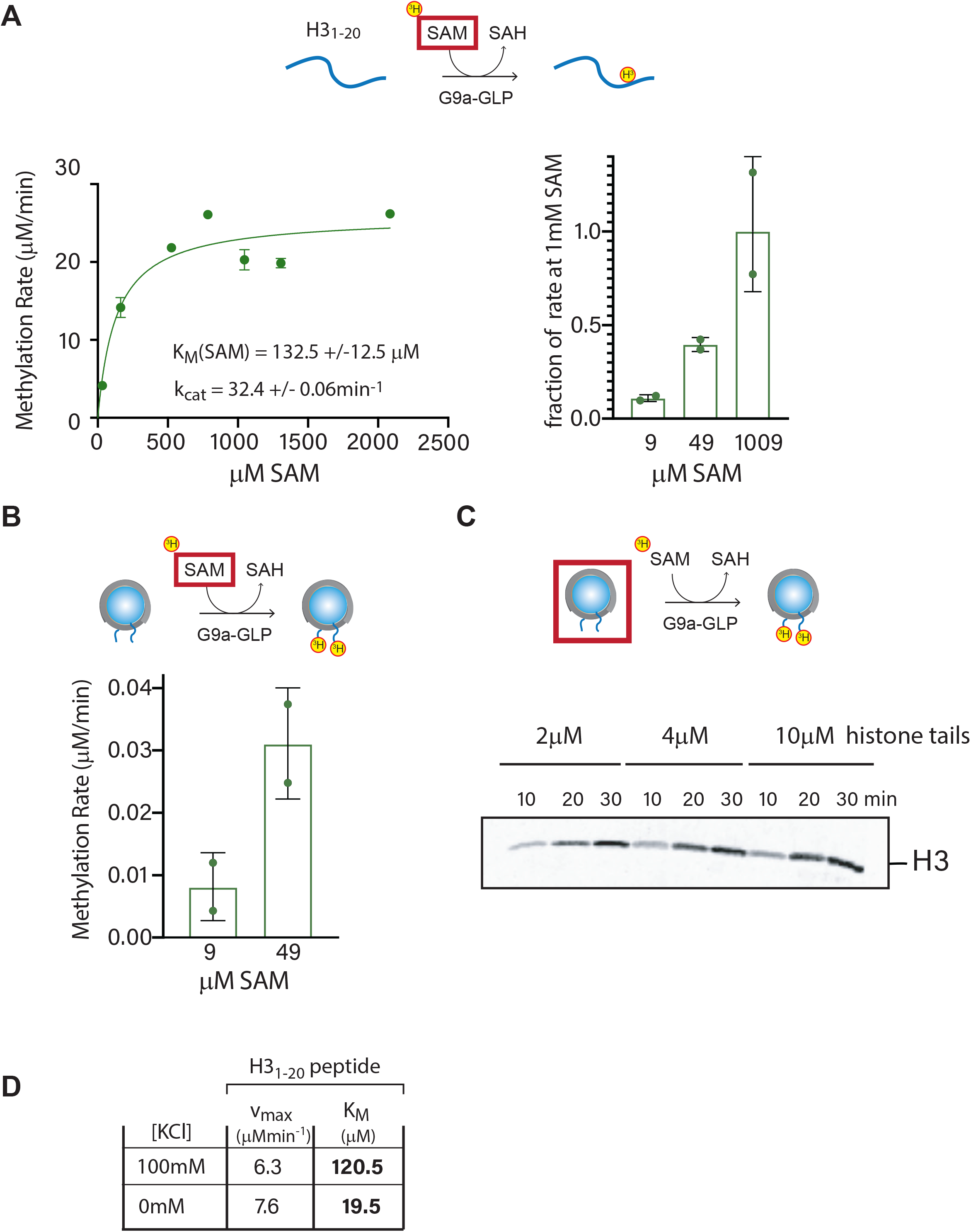
S-adenosyl methionine K_M_ and salt effects on the peptide K_M_. **A**. LEFT: The K_M_ for SAM was determined with G9a-GLP and H3_1-20_ peptide substrates. Initial rates are plotted at indicated total SAM concentrations and the curve fit to V = Vmax*[S]/(K_M_ + [S]). RIGHT: The initial rates of methylation by G9a-GLP on H3_1-20_ peptide were re-measured independently with different concentrations of cold SAM, while keeping ^3^H-SAM constant at 9μM. **B**. Initial rates of methylation by G9a-GLP on mononucloesomes (5μM) was measured with either only ^3^H-SAM (9μM) or 9μM ^3^H-SAM plus 40μM cold SAM. **C**. Example Tritium screen exposures used for the Michaelis-Menten curve in Figure 3 E (G9a-GLP). **D**. Kinetic parameters of G9a-GLP methylation of H3_1-20_ peptides under 100mM or 0mM (no salt) KCl. Note the drop in K_M_ under no salt. Error bars in A. and B. indicate standard deviation of two repeats.

**Supporting Figure 4:**
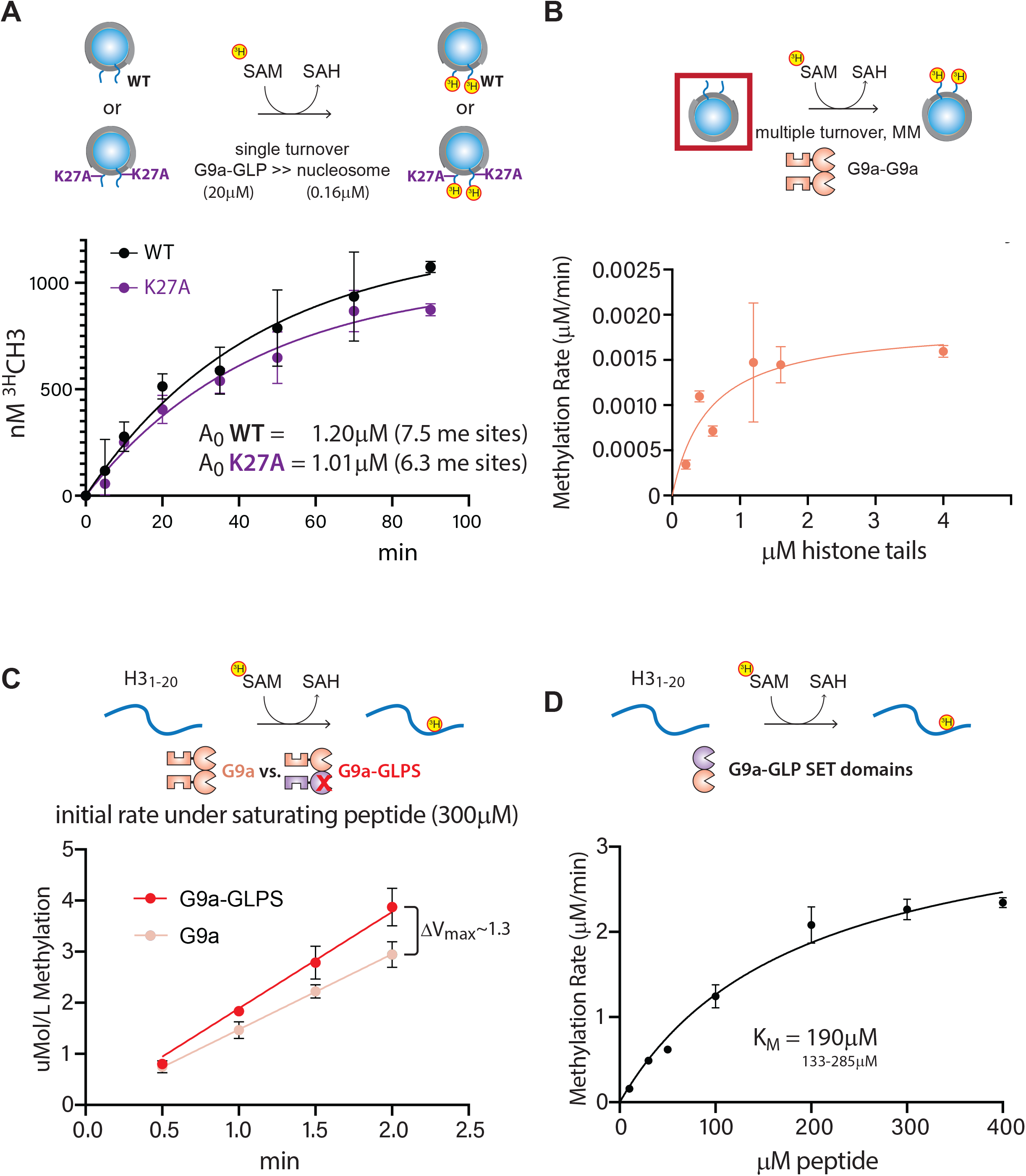
Role of K27 in nucleosome methylation by G9a-GLP. G9a-GLPS, G9a, and G9a-GLP SET domain methylation kinetics. **A**. Estimation of methylation at H3K9 vs. H3K27 by G9a-GLP on mononucleosomes. Methylation was conducted under Single Turnover conditions, with enzyme>>substrate (20μM G9a-GLP, 0.16μM WT or K27A mononucleosome). The methylation time course was fit to P(t)=A_0_*(1-e^- kt^), where k is the k_obs_ of the reaction. Solving for A_0_ yields the number of total methylation events, and given the known [mononucleosome], allows estimation of methylation events/nucleosome. **B**. A magnification of Michaels-Menten fit for G9a-G9a Multiple Turnover kinetics as shown in Figure 3E. **C**. Initial rate measurements for G9a and G9a-GLPS under saturating concentration (V_max_) of H3_1-20_ (300μM). The estimated maximal rate is only marginally different. **D**. Michaelis-Menten multiple turnover curve for G9a-GLP SET domains with H3_1-20_ peptides. The K_M_ is not lower than ANK-SET versions. All error bars indicate standard deviation of two replicates.

**Supporting Figure 5:**
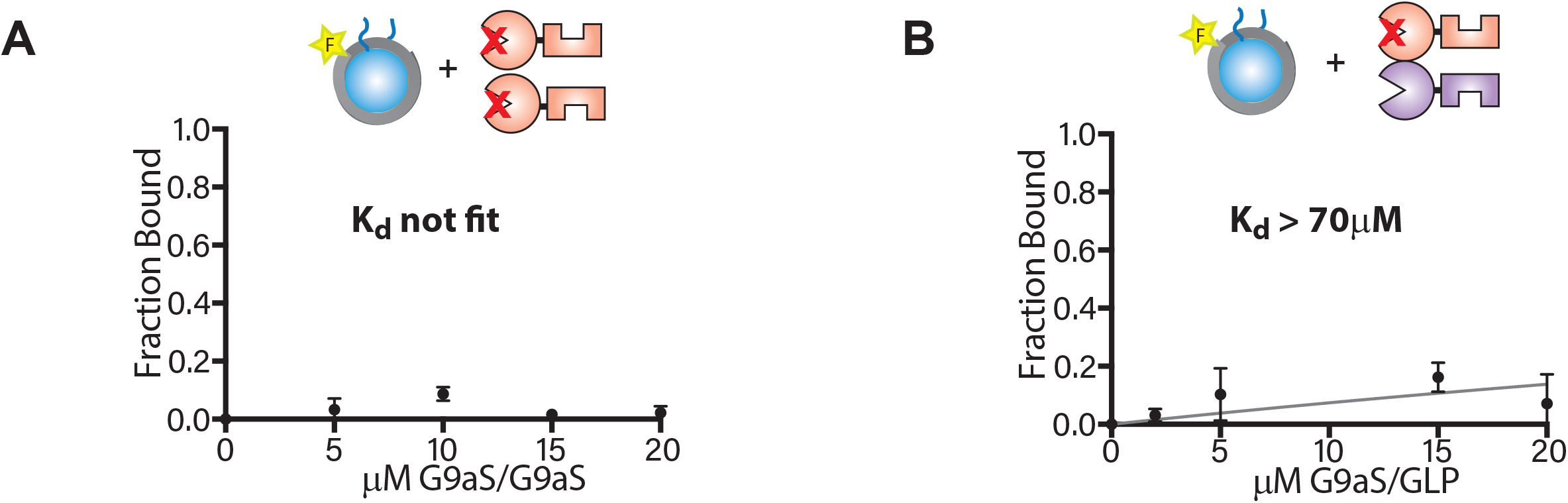
The SET domain active site of G9a is required for nucleosome binding. Fluorescence polarization of mononucleosomes containing 147bp 601 DNA 5′ fluorescein-labeled bound to **A**. catalytically inactive G9aS-G9aS and **B**. G9aS-GLP. Error bars indicate standard deviation of two replicates.

**Supporting Figure 6:**
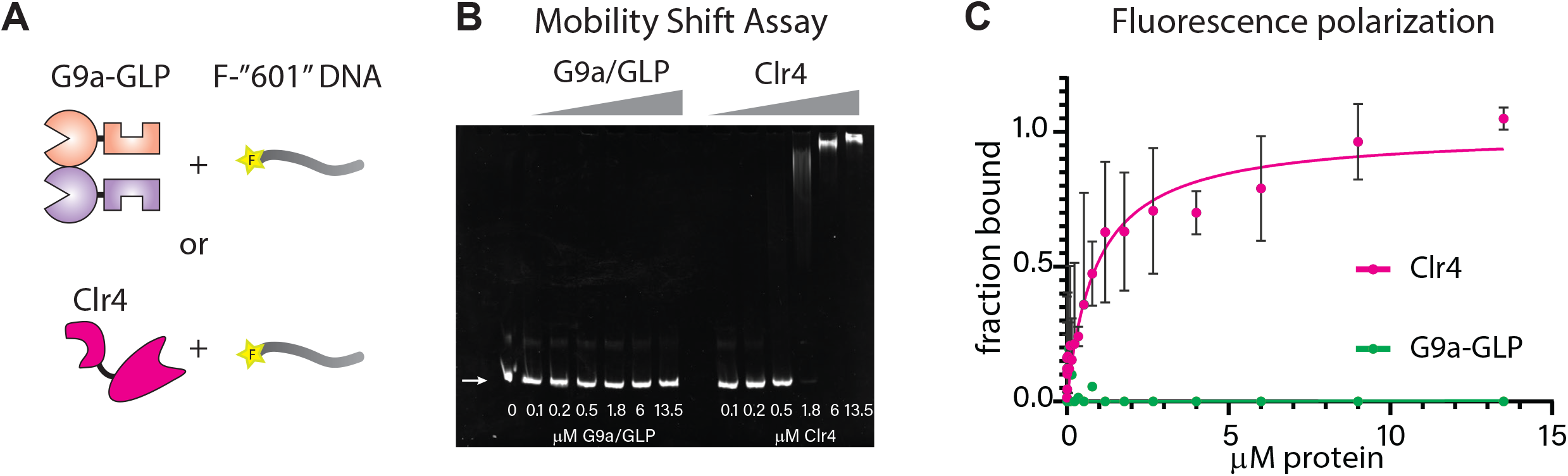
G9a-GLP does not bind DNA. **A**. G9a-GLP heterodimer or the monomeric fission yeast Clr4 H3K9 methyltransferase were assessed for binding the fluorescein-labeled 147bp ‘601’ mononucleosome DNA template. **B**. Electro-Mobility Shift Assay with G9a-GLP or Clr4 at indicated concentrations. **C**. Fluorescence polarization assay with G9a-GLP or Clr4 at indicated concentrations.

**Supporting Figure 7:**
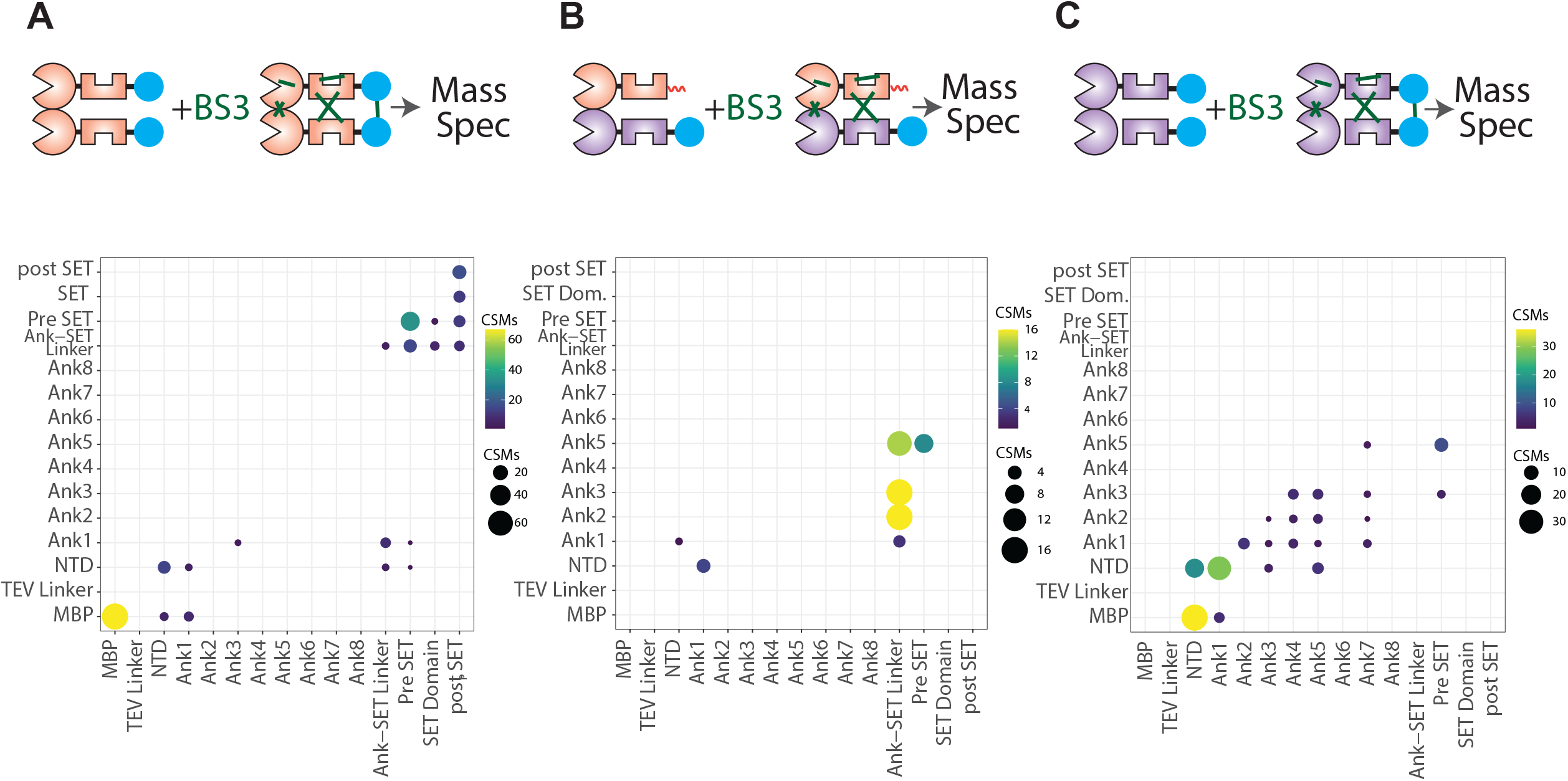
Crosslinking Mass Spectrometry for tagged G9a, GLP, and G9a-GLP. Crosslinking Mass Spectrometry for MBP:G9a (**A**), His:G9a-MBP:GLP (**B**), and MBP:GLP (**C**). Dot size and color indicates number of crosslinked spectral matches (CSM) summarized for domain pairs. Note that G9a and GLP data by necessity include intra-and inter protomer crosslinks. Note crosslinks involving the MBP tag.

**Supporting Figure 8:**
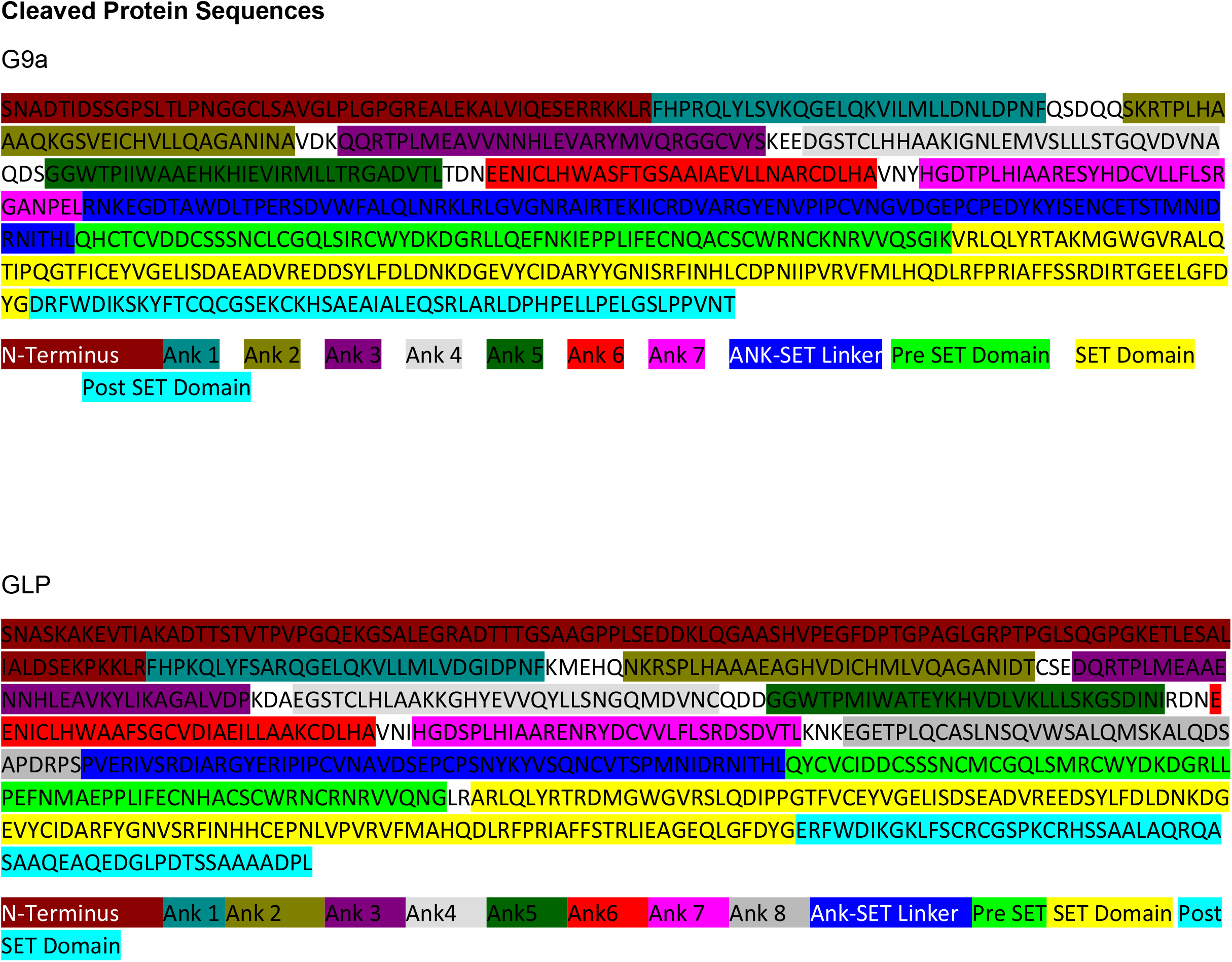
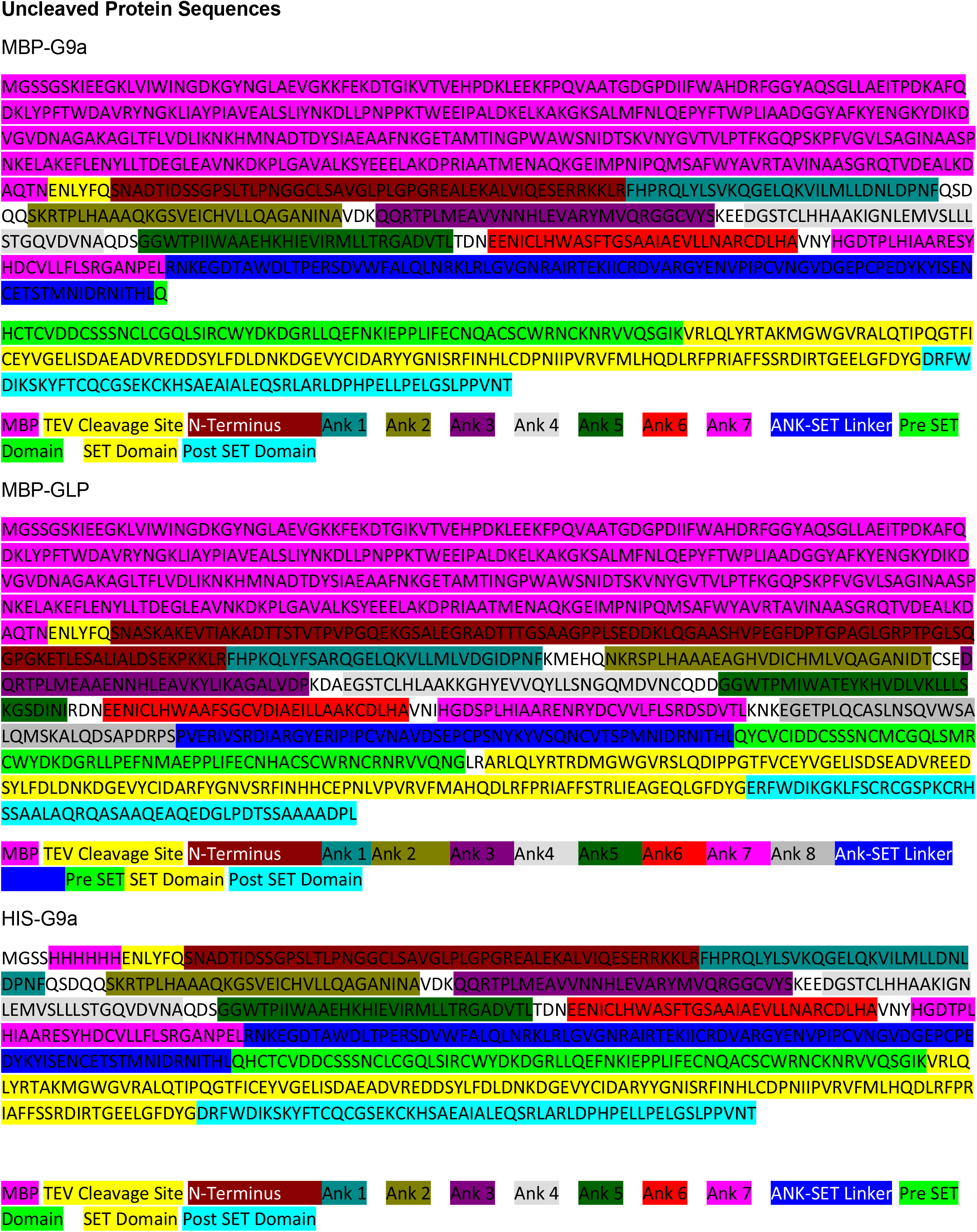
Domain annotations of tagged and tag-cleaved G9a and GLP proteins. Page 1: Amino acid sequence of the tag-cleaved G9a and GLP proteins. Domain identifiers are based on Uniprot and used in Figure 4 D-F and SFigure 7 A-C. Page 2: Annotated amino acid sequence of the tagged G9a and GLP proteins.

**Supporting Table 1: Crosslinking Mass Spectrometry data table**. Source data for the plots in Figure 4 D.- F. Q96KQ7, Uniport ID for G9a, Q9H9B1; Uniprot ID for GLP. Domain identifiers as in SFigure 8.

## Notes

### Competing Interest Statement

The authors have declared no competing interest.

### Summary of Updates

1.We repeated and expanded our binding measurements with higher [enzyme]. The new data significantly improves the quality of the data and solidifies our conclusions. In addition to the peptide data, we now prove binding measurements for all three enzymes to unmethylated, mono, and di- methylated nucleosomes. 2.For the methylation kinetic data, we have added new data comparing heterodimer to homodimers on nucleosomal arrays, as well as resolving whether the me0-me1 or me1-me2 transition is sensitive to acceleration by G9a-GLP on nucleosomes. 3.We additionally performed crosslinking- Mass Spectrometry studies on all three enzymes to test whether any internal protein contacts correlate with their particular behaviors. We find that the constitutive interactions between different domains and subdomains to be strikingly different between GLP, G9a, and G9a-GLP dimers, indicating different enzyme conformations despite the high homology between the G9a and GLP amino acid sequences. Surprisingly, we find interdomain contacts in the G9a-GLP heterodimer that not present in either homodimer.

